# Sequencing Metrics of Human Genomes Extracted from Single Cancer Cells Individually Isolated in a Valveless Microfluidic Device

**DOI:** 10.1101/258780

**Authors:** Rodolphe Marie, Marie Pødenphant, Kamila Koprowska, Loic Bærlocher, Roland C.M. Vulders, Jennifer Wilding, Neil Ashley, Simon J. McGowan, Dianne van Strijp, Freek van Hemert, Tom Olesen, Niels Agersnap, Brian Bilenberg, Celine Sabatel, Julien Schira, Anders Kristensen, Walter Bodmer, Pieter J. van der Zaag, Kalim U. Mir

## Abstract

Sequencing the genomes of individual cells enables the direct determination of genetic heterogeneity amongst cells within a population. We have developed an injection-moulded valveless microfluidic device in which single cells from colorectal cell (LS174T, LS180 and RKO) lines and fresh colorectal cancers are individually trapped, their genomes extracted and prepared for sequencing, using multiple displacement amplification (MDA). Ninety nine percent of the DNA sequences obtained mapped to a reference human genome, indicating that there was effectively no contamination of these samples from non-human sources. In addition, most of the reads are correctly paired, with a low percentage of singletons (0.17 ± 0.06 %) and we obtain genome coverages approaching 90%. To achieve this high quality, our device design and process shows that amplification can be conducted in microliter volumes as long as extraction is in sub-nanoliter volumes. Our data also demonstrates that high quality single cell sequencing can be achieved using a relatively simple, inexpensive and scalable device.

## Introduction

Standard molecular methods that analyse DNA sequences in populations of cells need sufficiently deep sequencing to detect heterogeneity at any given location of the genome and do not clearly define the co-occurrence of mutations in a given cell, and so do not generally define the genetic heterogeneity in a tissue, especially cancers.

Cancers arise from a somatic evolutionary process in which mutations, or relatively stable epigenetic changes, are successively selected for and therefore occur with relatively high frequency in the population of cancer cells. It is these genetic and epigenetic changes that determine the properties of a cancer and so are the major determinants of prognosis and of the responses to different treatments. With the extraordinary development of DNA sequencing technology, there is now extensive data on the types and frequencies of the major, so called ‘driver’, mutations found in a wide variety of cancers, and in some cases very clear evidence of the relationship between the mutational content of a cancer, and its response to therapy. As a result, there is now great interest in DNA sequencing of single cancer cells to determine the nature and extent of clonal genetic heterogeneity in a given cancer.

Treatment can then be directed at the different clones that co-exist in the cancer and thus single cell DNA sequencing (1) becomes an extremely important tool for matching the treatment of a cancer to its genetic make up. This is the essence of precision medicine as applied to cancer treatment. There is also an interest in single cell mRNA analysis (2), which can help to identify gene expression differences, due mainly to DNA methylation as well as interest in examining methylation directly (3). There has therefore been increasing focus on the development of methods that obtain molecular information from single cells by isolating and sequencing their DNA and mRNA content (4–6). Similarly, in metagenomics, where bacteria, fungi and other microbes may not be culturable, a robust single cell analysis is important to evaluate the genomes of the distinct microbes present in a sample (7). In addition to untangling heterogeneity, the single cell methods are relevant to cases when only a small number of cells is available, for example in the analysis of circulating tumour cells (8,9) and circulating fetal cells in maternal blood (10).

A single diploid human cell contains around 7 picograms of genomic DNA and some form of amplification is therefore needed to obtain the amounts of material necessary for current sequencing methods. Amplification by multiple displacement amplification (MDA) (11) and PCR based methods such as DOP-PCR (12), Picoplex (13) and, Multiple Annealing and Looping-based Amplification Cycles (MALBAC) (14) have been used relatively effectively to do single cell whole genome DNA sequencing. Genome coverage of >90% has been claimed to be routinely obtained from single cells using MDA and MALBAC, with a MDA kit adapted for single cells reportedly giving superior all round performance (12). However, issues around uniformity of coverage, allelic dropout, false positives (amplification and/or sequencing errors) and unmappable reads (e.g. from primer-dimers), remain (15).

The MDA process results in uneven coverage across the genome. Some of the amplification biases are presumed to be a result of stochastic effects due to the sampling of a small number of molecules. In MALBAC amplification bias can be corrected by normalizing the GC content (16). Another approach to dealing with this problem is the use of barcoding or identification tags (17–19).

Contamination, if not carefully controlled can lead to difficulty in interpreting results and limits the sequencing capacity that is available for a single cell of interest. To combat this, single cell genomics is preferably conducted in a clean room (15), in microwells (20) or in a microfluidic device (7,11,16). Contamination can also be assessed by the use of appropriate known genetic markers.

Existing microfluidic devices (7,16) requiring multiple-PDMS layers (21), one layer for the passage of fluids and another for valves to control the fluids through the device, are difficult and expensive to manufacture because PDMS casting is not a scalable industrial process. We introduce a novel valve-less microfluidic device for single cell genomics that is manufactured in a thermoplastic material by injection moulding, a process that is scalable at low cost (22). Our chip design is based on a hydrodynamic cell trap (23) derived from a previously described device for cell culture (24). Our device design enables the process to be carried out on an optical microscope or in “Cell-O-Matic” a specially built single cell processing instrument (Philips BioCell), in either case allowing us to monitor both cell trapping and genome extraction.

Our single cell sequencing data obtained using the Cell-O-Matic instrument show that we can achieve reasonable levels of whole genome coverage in a significant proportion of cells. Our results compare well with other reported whole genome sequencing from single cells using instrumentation in which the amplification is performed in nanoliter-reaction chambers. In our device only the DNA extraction occurs in a sub-nanoliter volume of solution, while the amplification is performed by adding microliter volumes of reagents in the device outlet. We conclude that the critical step in single cell whole genome amplification with regard to sequence allelic dropout, contamination and genome coverage is to elute/extract DNA in sub-nanoliter volumes in the confinement of the microfluidic device, while performing the amplification in such small volumes may only be required for high reduction of reagent consumption, but at the cost of higher device complexity and cost.

## Material and Methods

### Device Fabrication

The device is fabricated by injection moulding of TOPAS 5013, a Cyclic Olefin Copolymer with a glass temperature of 130°C as described elsewhere (25) but with some modifications. First, the microchannel network is defined to have a depth of 30 μm in a silicon substrate by UV lithography and reactive ion etching. All dimensions of the microchannel network can be found in Supplementary Figure S1A-F. Next, a Ni/V seed layer is deposited on the silicon master and Nickel is electroplated at a final thickness of 300 μm. Silicon is removed by KOH etching.

The Nickel shim is cut to fit in the mould of the injection moulder (Engel, Germany). The mould creates a 2 mm-thick, 50 mm diameter disc replica of the shim and 12 holes through the disc used to connect the microfluidics. It also creates for each hole, a female LUER connector on the side opposite the microfluidics side. The temperature of the mould is regulated such that injection moulding is performed using a variotherm process (26) allowing the polymer part to be moulded and removed from the mould at different temperatures. The shim is replicated using a mould temperature of 115°C, a shim temperature of 155°C, a holding pressure of 1150 bar and a cooling time of 14 seconds so the demoulding temperature is below 126°C.

The microfluidics chips are sealed with a 150 μm-thick TOPAS 5015 foil using UV assisted thermal bonding. The initial roughness of the foil is reduced by a hot embossing process on a hydraulic press (P/O/Weber, Germany) at 140°C and 5.1 MPa using two smooth Nickel discs electroplated from silicon wafers. The glass transition temperature (Tg) of TOPAS 5013 is 130°C. The devices and lids are exposed to UV-light from a mercury arc lamp for 30 s and then placed in an Aluminium holder that fits to the LUER fittings. A smooth nickel disc and a thin PDMS plate were placed on top of the lid to ensure a uniform pressure across the device during bonding at 125°C and 1.5 MPa for 3 minutes.

### Instrument

The microfluidic device can either be mounted on a conventional epi-fluorescence microscope or on a custom designed instrument. The instrument is a modified Philips BioCell Fluidscope (6) (Supplementary Figure S1G). In brief, the single use microfluidic chip is mounted on a stage equipped with translation (y-axis) as well as temperature control via three Peltier elements. The temperature inside the wells of the chip is calibrated and regulated by a proportional–integral–derivative (PID) controller using the temperature of the stage as input. The lid closing the microfluidic chip wells is connected to a multi-channel pressure controller (MFCS-EZ, 300 mbar range, Fluigent). The chip is imaged using an objective (10x, NA 0.45 Wild Heerbruug) mounted on a translation stage (x-axis) and focusing by z-translation. Epi-fluorescence imaging is performed with an excitation at 470 nm (LED, Thorlabs model M470L3) equipped with collimator lenses. A dual band filter cube (Semrock, excitation filter model 733-495/605-Di01-25x36, dichroic mirror model 733-474/23-25, emission filter model 733-527/645-25) allows imaging of green fluorescence and bright field. The bright-field illuminator delivers light from the top side of the device through a window in the lid using a LED with center wavelength of 505 nm and a 20 nm bandwidth. A CMOS imaging sensor (Fairchild, CIS1910) with 1920 x 1080 pixels, and pixel size 6.5 μm, is used with a home-built image board. Finally, all the elements are controlled through software which guides the user through a pre-established workflow for priming, cell capture, lysis and amplification.

### Device Operation

(see supplementary Protocol S8 for the detailed operation procedure)

Solutions are pipetted into the LUER connectors used as reservoirs containing up to 50 μL of solution. Air pressure supplied by the pressure controller attached to the wells drives the flow inside the device. We apply pressure to the cell inlet as well as the buffer inlets B1 and B2 (Figure 1) while the waste and the trap outlets are always left at atmospheric pressure. For cell capture, pressures in the range of 5 to 10 mbar were applied, corresponding to sample flow rates of 2.9 - 5.7 μL/h. Details of the pressure settings used are given in the supplementary Protocol S8.

**Figure 1.**
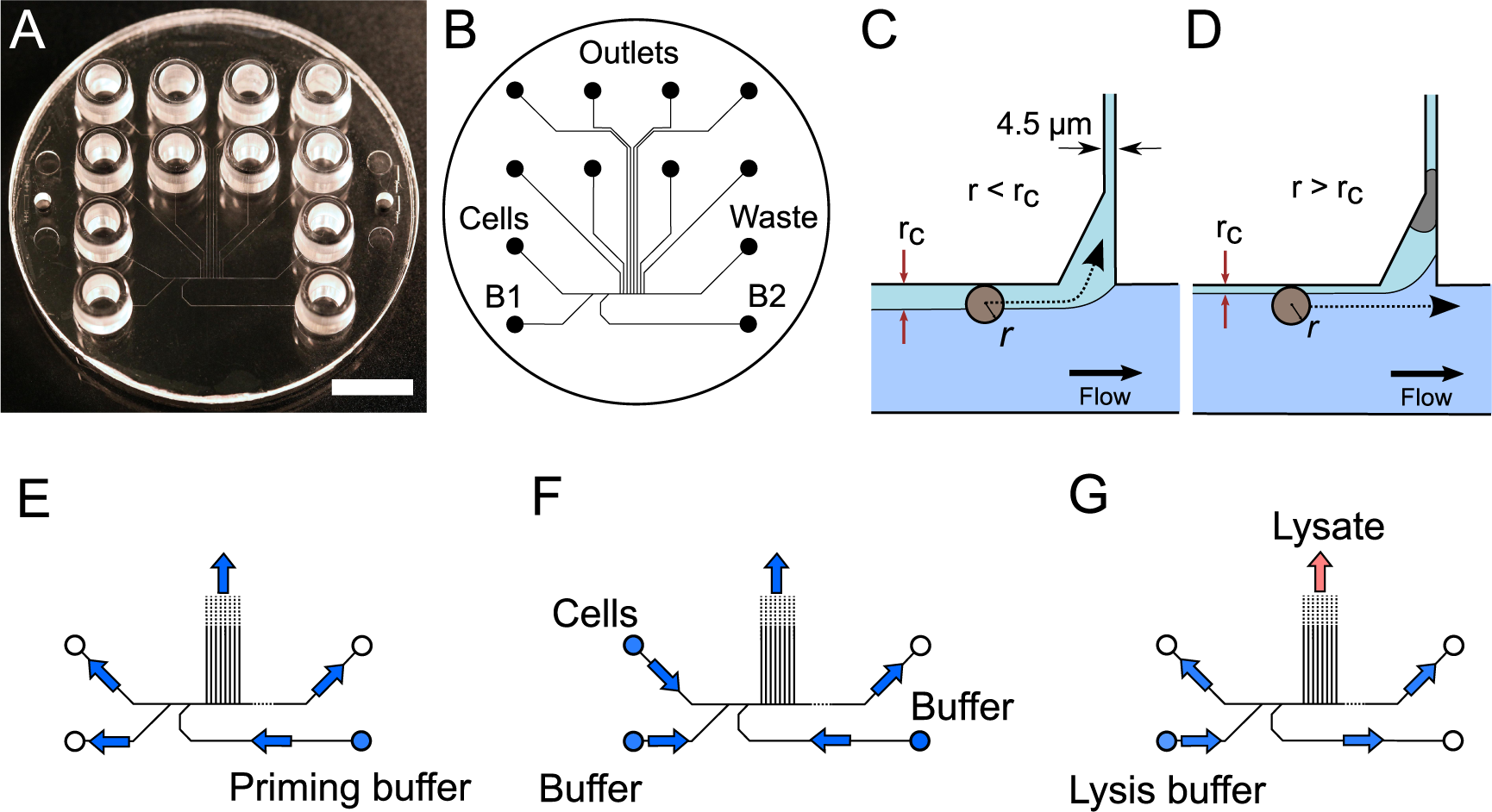
Device design and operation. A) Image of the single use polymer device. Scale bar is 1cm. B) Microfluidics layout. C) Conditions for cell trapping an unoccupied trap *r<r_c_*. D) The trapped cell reduces the flow through the trap such that for the next incoming cell, r>r_c_. E-G) Flow directions in the device under priming, cell capture and lysis.

### Wet lab

Cells from colorectal cancer cell lines LS174T, LS180 or RKO (concentration: 6×10^5^ cells/mL) were stained with 1 mM calcein AM and suspended in BD FACSFlow buffer. After cell capture, the trap occupancy is checked by bright field and fluorescence imaging of the calcein signal. After trapping cells, the B1 and B2 inlets were emptied leaving negligible volumes in the outlets.

### On-chip protocol 1: (39 cells, processed in Oxford)

The devices were primed with degassed 0.1% v/v Triton X-100 in BD FACSFlow buffer (1 minute at 200 mbar) followed by degassed 0.1 mg/mL BSA in BD FACSFlow buffered (2 minutes at 200 mbar).

The cell suspensions were loaded onto the chip and cells were trapped (see supplementary Protocol S8).

The alkaline lysis buffer of a commercial MDA kit was used to lyse the trapped cells (REPLI-g UltraFast Mini Kit, Qiagen). The lysates were pushed with 20 μl of the alkaline lysis buffer, incubated for 20 minutes at room temperature, and then pushed with 20 μl of the neutralisation buffer for 20 minutes at room temperature. (For further details see supplementary Protocol S8).

### On chip protocol 2: (13 cells, processed in Eindhoven)

In a second experiment, we process cells in a different laboratory using a different cell lysis based on proteolysis. The protocol is essentially the same as protocol 1 with a few modifications. The device was primed with ethanol from all inlets (cells, B1 and B2) then BD FACSFlow buffer was loaded in all wells before cells were loaded. In all the samples, the cell cytosol is removed by a solution of 0.5% triton-X100 in 0.5xTBE buffer, containing 1 μM of YOYO-1 dye (6). The DNA was eluted from the cell traps by proteolysis buffer (13 cells) containing >200 μg/mL protease K, 0.5% v/v Triton-X100 in 0.5xTBE buffer. Then alkaline lysis solution from the REPLI-g Single cell kit was added to the outlet well (6).

### Whole genome amplification in device outlets

Whole Genome Amplification (WGA) was performed using REPLI-g Single cell or the Qiagen REPLI-g UltraFast Mini Kit (Qiagen) which are based on Multiple Displacement Amplification (MDA) technology to amplify gDNA from small samples. 10 μL of reaction mix with Phi29 polymerase was added to the chip outlets containing gDNA. The device was left on the heated stage of the instrument (alternatively placed in a thermal cycler) for 8 hours at 30°C. The final step was 5 min at 65°C to inactivate the polymerase. All WGA steps were done on chip. Following this the amplified DNA was transferred to PCR tubes, tested for quality and, subject to passing quality control, sent to Fasteris for library preparation and DNA sequencing.

### Multiplex chromosome check at Oxford (off-chip)

To confirm the quality of WGA products, multiplex PCR was performed. (See http://www.sigmaaldrich.com/technical-documents/articles/life-science-innovations/qualitative-multiplex.html). Five primers targeting 5 chromosomes (2, 4, 12, 13 and 22) were used. PCR products (295 bp Chr13, 235 bp Chr12, 196 bp Chr2, 150 bp Chr22 and 132 bp Chr4) were visualized on a 4% agarose gel (see supplementary Protocol S8). Those 5 positions were chosen based on https://www.sigmaaldrich.com/technical-documents/articles/biology/ffpe-wga-poster.html.

### Library preparation and sequencing

Samples for sequencing were quantified using a Qubit fluorometer instrument before starting the library preparation procedure. Depending on the initial sample concentration, the library preparation was done using the Illumina Nextera or the Illumina Nextera XT kits. Sequencing reactions were performed for all samples using Illumina HiSeq technology as 2x125 base pairs high-output runs.

### Bioinformatics

All libraries sequenced were then mapped against the human genome GRCh37.

We report on the percentage of reads mapped to the human genome (Figure 2A) and the total number of reads (Figure 2B). We also use the cumulative fraction of bases covered to generate Lorenz plots (Figure 3) and the coverage plots (Figure 4).

**Figure 2.**
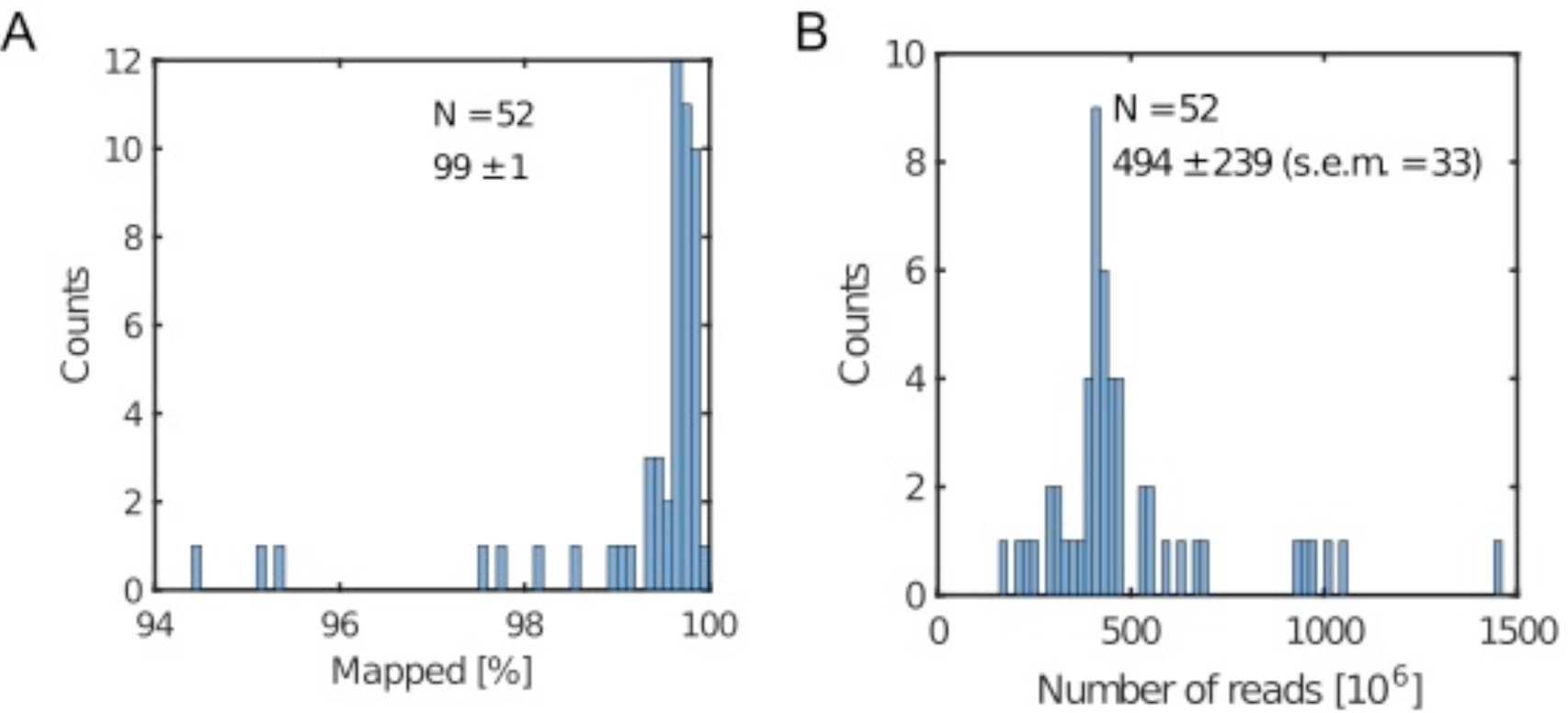
Metrics. A) Percentage of mapped reads B) Total number of reads. Legend displays the mean, the standard deviation and the standard error of the mean (s.e.m.).

**Figure 3.**
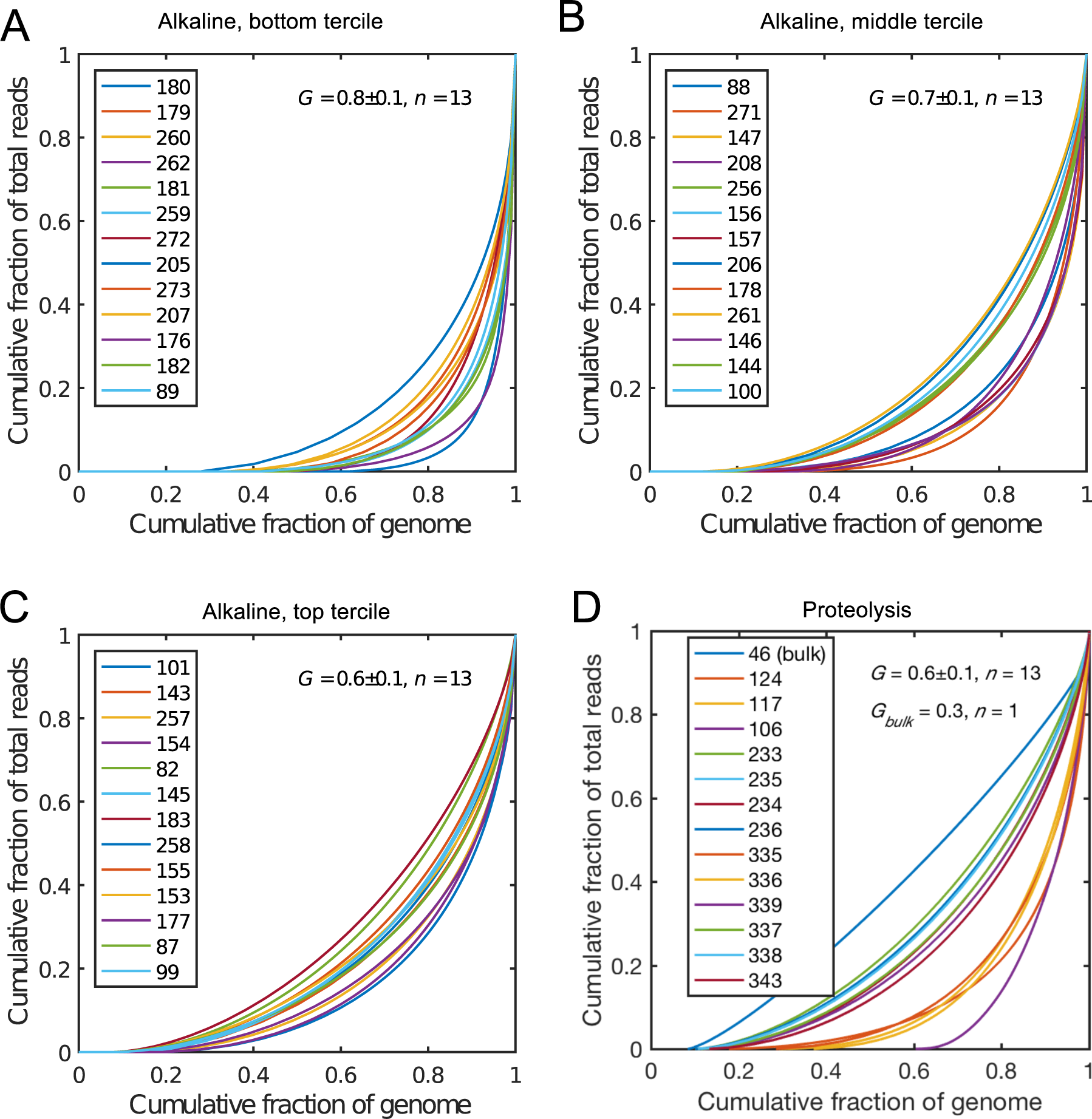
Lorenz plots for the cells processed by alkaline lysis in Oxford, in three groups of n = 13 cells: (A) bottom, (B) middle and (C) top tercile according to the percentage of non-covered bases in the genome. (D) Lorenz plot for the single cells processed by proteolysis in Eindhoven, n = 13 cells. Cells 124 to 236 are LS174T cells. Cells 335 to 343 are RKO cells. The bulk of LS174T is also shown (sample ID 46). We display the Gini coefficient *G* mean value and the standard deviation for each group.

**Figure 4.**
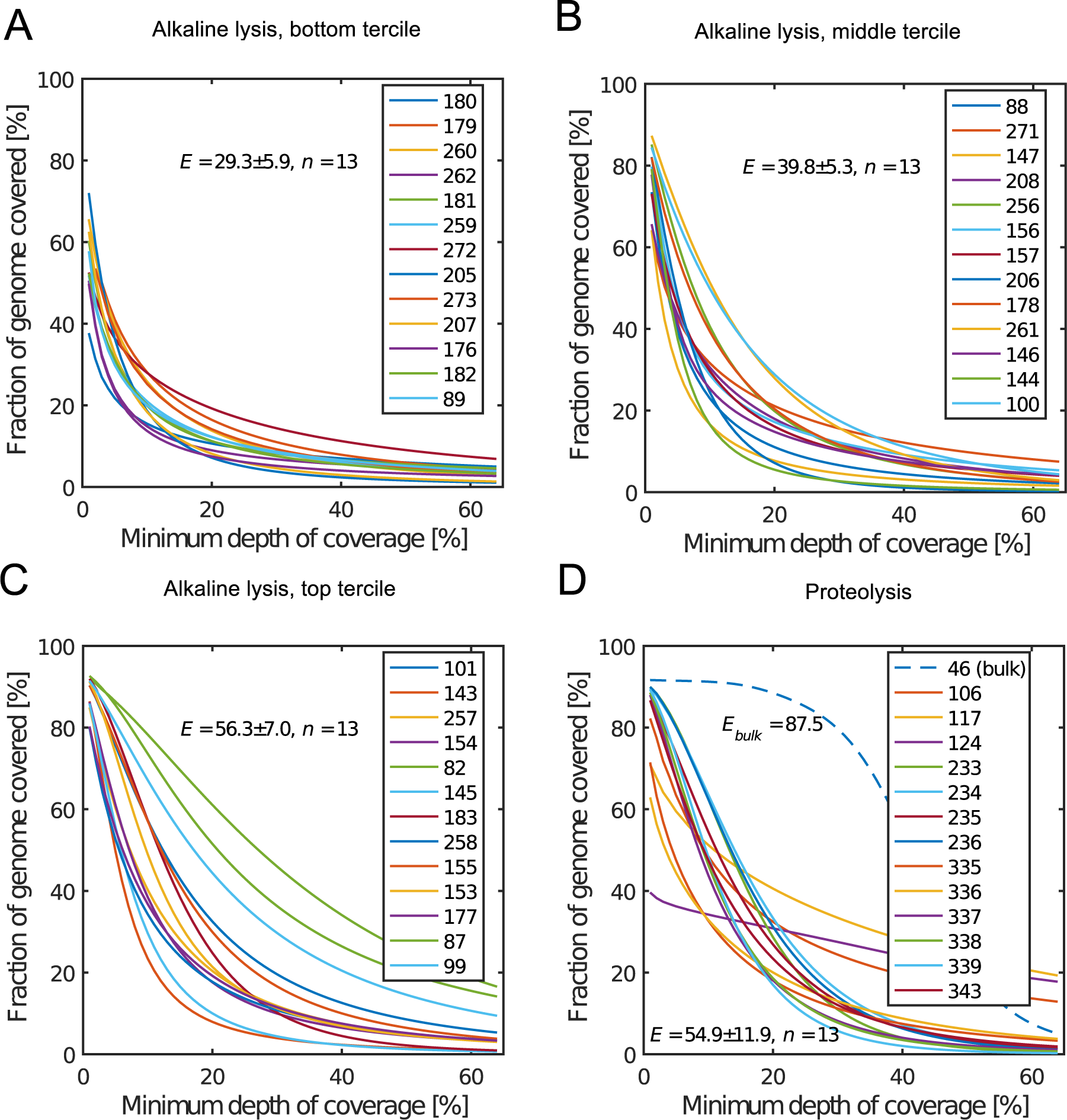
Coverage plots corresponding to the (A) bottom, (B) middle and (C) top tercile and (D) the single cells processed by proteolysis in Eindhoven. Cells 124 to 236 are LS174T cells. Cells 335 to 343 are RKO cells. The bulk of LS174T is also shown (sample ID 46). We display the E-score as mean value and the standard deviation for each group. The E-score is calculated from a normalized coverage curve as described in (Mokri et al).

For all samples, allelic dropout estimates (Figures 5 and 6) were based on a set of validated heterozygous variants, selected for each cell line: 12 SNPs were used for the LS174T and LS180 cells, 13 for the RKO cells (see supplementary Table S10). The known variants were searched for in the raw variant results, namely without applying any filter on the variant quality. All selected variants were correctly detected in the respective bulk samples as heterozygous variants. For single cells, a drop of a heterozygous variant to a homozygous call would reflect dropout of a single allele, while absence of any call would represent a dropout of both alleles (resulting in no coverage of the position). The procedure used for generating Figure 5 and 6 is described in detail in the results section.

**Figure 5.**
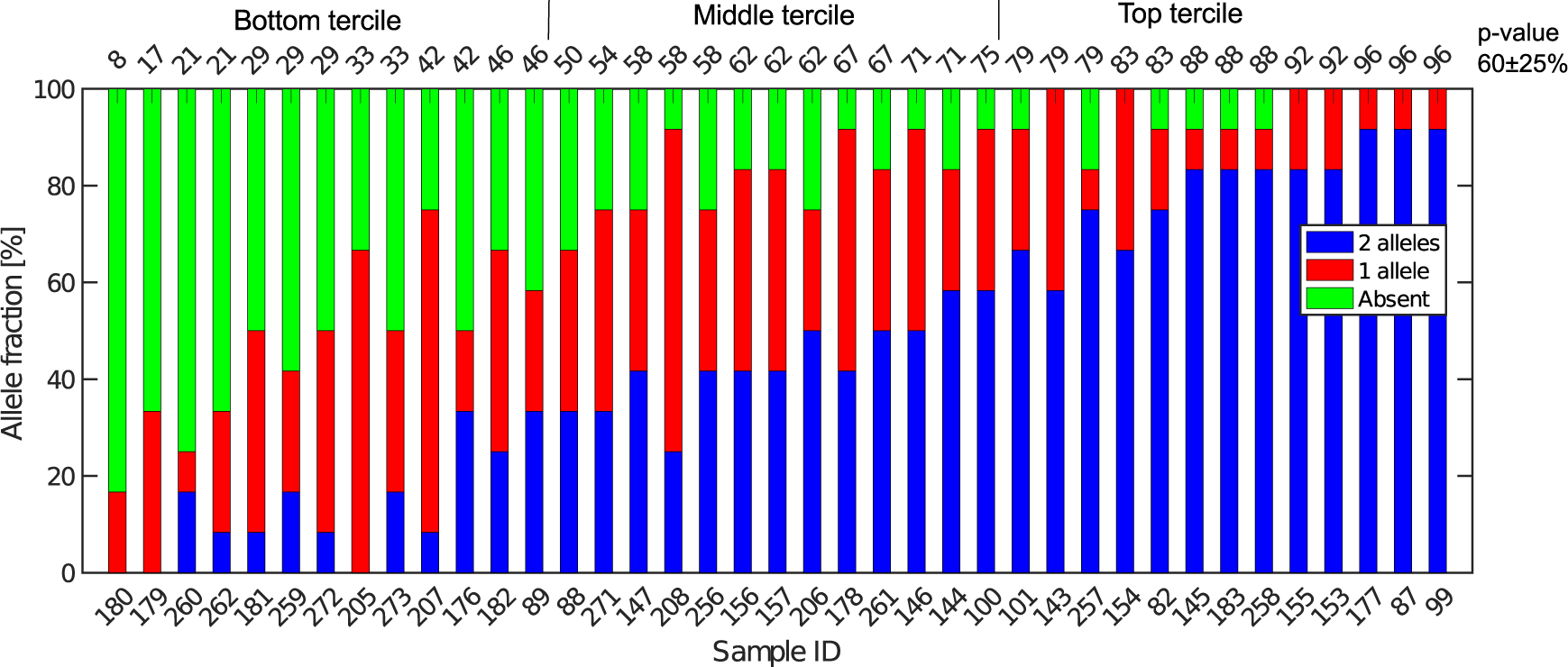
Extent of Allelic Dropout for 12 heterozygous SNPs selected for LS174T (36 cells) and LS180 cells (3 cells). On the x-axis, the sample ID for the cells analyzed. Cells are ordered by increasing p-value (in percent). The mean (SD) for all 39 cells processed in Oxford is 60 (25).

**Figure 6.**
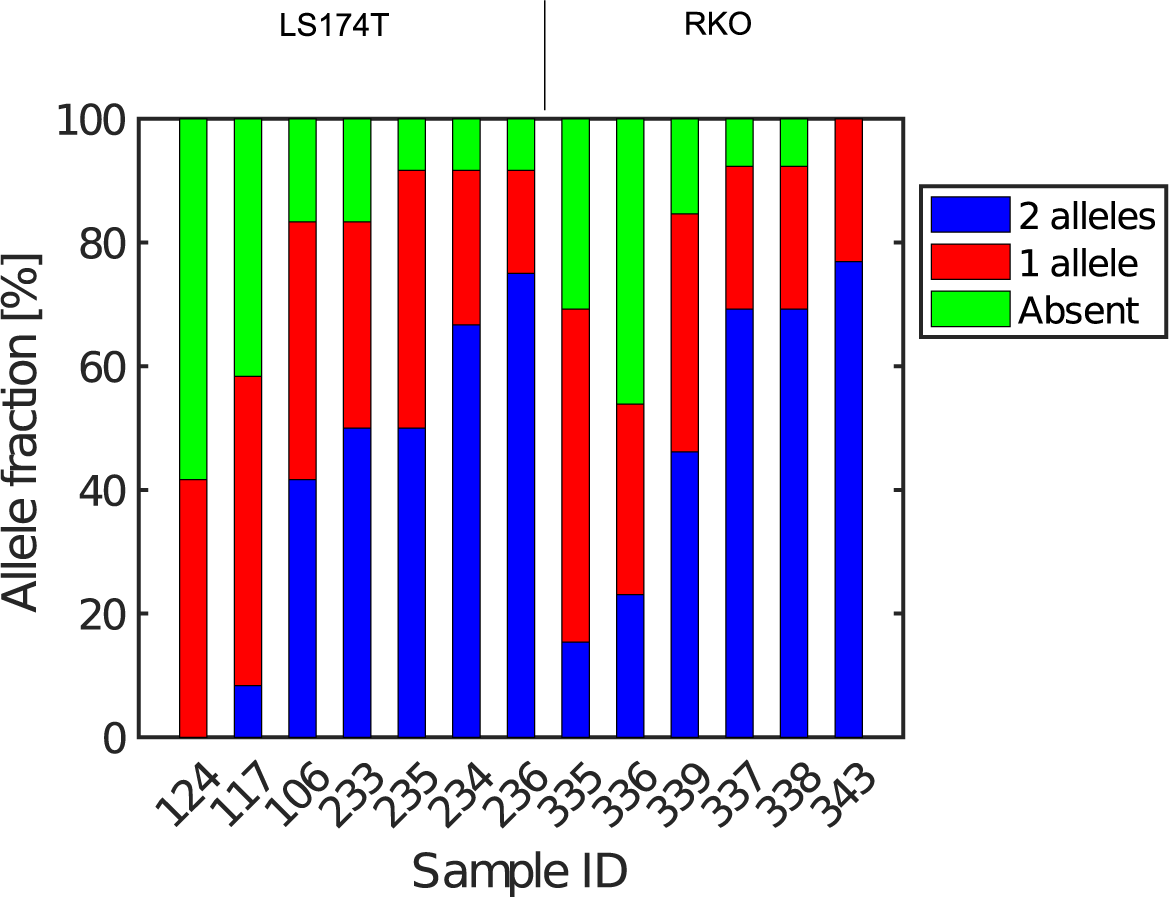
Extent of Allelic Dropout for heterozygous SNPs for cells processed in Eindhoven. Cells 124 to 236 are LS174T cells (12 heterozygous SNPs) and cells 335 to 343 are RKO cells (13 heterozygous SNPs).

From the Lorenz and the coverage plot we calculate the Gini coefficient *G* and the Eveness score *E* respectively. For the latter, the normalized coverage curve is used and scaled by the maximum fraction of genome covered in a bulk sequencing run of 10/9.136. This is done to account for the fact that in the bulk sequencing of LS174T, 8.36% of the bases remain uncovered (6).

## Results

We mainly used the LS147T cell line derived from a colorectal cancer (CRC) as a model system for singe cell analysis. This is a very well characterized cell line (see e.g. (27)) that has been widely used for CRC cell characterization and drug response studies (see e.g. (28,29)). In some experiments, we also processed cells from the RKO and LS180 cell lines (the latter was alternatively derived from the same CRC as LS174T), and from cells obtained directly from fresh CRCs.

For each experiment a single use microfluidic device (Figure 1A and supplementary Figure S2) is placed in an instrument derived from a Philips BioCell Fluidscope allowing bright field and fluorescence imaging, controlling the device temperature and connecting the device inlets (Cell, B1 and B2 inlet, Figure 1B) to a multi-channel air pressure controller (6) (supplementary Figure S2G). Fluorescence imaging and the use of YOYO-1 intercalating DNA dye enables monitoring of cell lysis. However, the dye may be omitted to avoid interference with the subsequent quality of the preparations with respect to their use for DNA, or RNA sequencing (6).

### Device design and fabrication

The microfluidic device is a single use passive device fabricated by injection moulding using a mould that creates 12 connectors placed at distances corresponding to a 96-well plate standard and can contain up to 50 μL of a solution (7) (Figure 1B). The microfluidic device has a single depth microchannel network with dimensions such that LS174T and similar colorectal epithelial cells are successfully transported and trapped (supplementary Figure S3). LS174T cells have a median diameter of 14 μm, which is in the characteristic size range for colorectal cancer cells, and so we designed the microfluidic network to be 30 μm-deep and at least 30 μm-wide except for the flow constrictions that are used as cell traps.

Our design is the result of iterative optimization where we identified and improved three critical aspects of the device design and fabrication: i) the flow through the trap, which depends on its cross section and the flow resistance of the outlet channels, ii) the shape of the cell pocket and iii) the moulding quality of the cell inlet.

The trap cross section has to be smaller than the LS174T cells size to retain the cells, but also sufficiently large so that it collects a significant fraction of the main flow in the feeding channel for cells to be directed through the trap. As a boundary condition, our choice of a single depth design means that the trap depth remains the same throughout the chip, namely 30 μm. As a result, the trap has a high aspect ratio, within the limit achievable during the fabrication of the master in silicon by micromachining. Finally, the fabrication by polymer replication results in the channels and in particular the trap having tilted sidewalls (up to 3 degrees) to allow the separation of the polymer part from the mould during injection moulding. As a result, the cell traps have a cross section 30 μm-deep, 4.5 μm wide at the bottom and 7.5 μm wide at the top.

The pocket receiving the cell has an asymmetric design (Figure 1C-D). This is in contrast to previously reported devices based on hydrodynamic trapping where flow focusing is used to direct the cell to a microfluidic constriction that is a bypass in an otherwise symmetric flow profile. In our device, the flow focusing is asymmetric since cells are aligned against the wall of the feeding channel. A symmetric pocket creates a dead volume after the constriction (Supplementary Figure S3) that is a spot where a cell decelerates and can settle just outside the cell trap. By making the pocket asymmetrical, we improve the flow profile such that cell trapping is more efficient. The optimized design gave the best results in terms of numbers of traps per chip having single cells.

Finally, the connection of the feeding channel and the well receiving the cells is a critical aspect of the design. The surface roughness at the inlet is of paramount importance since a sharp edge tends to stop cells entering the channel. The injection moulding parameters are therefore adjusted to produce a round edge. In addition, we ensured that the shim is mounted into the injection moulder only once. This greatly improved the quality of the final device since successive mounting of the shim increases the roughness at the connections with the inlets due to the alignment tolerance of the shim in the mould.

On the optimized device (Figure 1), single cells were trapped routinely, on average, in 3 to 6 out of 8 possible traps. On some occasions, cell doublets are trapped and this may be because cell doublets enter the device in the first place. For that reason, cell traps are imaged in bright field and fluorescence after trapping to confirm the presence of, and then exclude, such cell doublets from further analysis.

### Cell capture and lysis

For each experiment, solutions are loaded in the device wells and a single pressure is applied to the inlets either to wet the device, trap cells or lyse the trapped cells (Figure 1E-G and supplementary Protocol S8). After priming of the microfluidics, cells loaded in a cell inlet are pushed into the feeding channel where they become aligned against a sidewall of the channel by the incoming flow from the buffer inlets (B1 and B2 in Figure 1B), similarly to a recently described cell-trapping device (30). Once aligned, a cell of radius *r* follows a streamline at a distance *r* from the sidewall. The cell traps are constrictions that connect the feeding channel with separate outlets. A trap collects a fraction of the flow from the feeding channel such that we call *r_c_* the position of the last streamline entering the trap. A cell of radius *r* enters the trap downstream if *r<r_c_* (Figure 1C). Since a trap is only 4.5 μm-wide, a cell entering it is captured in the constriction. This cell then occupies a pocket recessed from the main flow through the feeding channel. A cell cannot block the flow through the trap completely since the channel depth (30 μm) is much larger than the cell. However, the flow resistance is sufficiently increased for the next incoming cells to pass by and be directed to the following free trap (Figure 1D). Cells that are not trapped are collected in the waste outlet. It is possible routinely to achieve 3-6 single cells in the 8 possible traps.

Cellular DNA is eluted from the cell trap by introducing a lysis solution from the inlet B1 (Figure 1 B).In our study, we compared two lysis solutions. For one, a solution for proteolysis including proteinase K and Triton-X100 was used for the 13 cells (LS174T and RKO) whose results were obtained in Eindhoven. This lysis solution enables collecting the RNA prior to collecting the DNA of the trapped cell (6). Alternatively, an alkaline lysis buffer (D2, pH above 12) provided with the Repli-g UltraFast kit was used in Oxford for the analysis of the 39 single cells from the LS174T and LS180 cell lines, and from two fresh tumour samples (see supplementary Figure S3). The alkaline lysis is the one adopted in commercially available kits for eluting DNA for sequencing. Both solutions successfully lyse the cells trapped and elute the DNA from the trap as observed in experiments where the DNA is labelled with an intercalating dye so it can be visualized by fluorescence microscopy. From the results of the single cell sequencing using the two different approaches, as discussed below, we conclude that both approaches to lysis were appropriate for MDA. This is, perhaps, surprising in the case of DNA extraction by proteolysis since proteinase K might be expected to digest the polymerase. However, there are six orders of magnitude difference between the volumes of lysate (pL) and the volumes of the reagents added to the well (μL), which thus makes the protease content in the MDA mix insignificant (6). The success of the amplification and sequencing is the best indication that the lysis is successful.

DNA samples that successfully amplified were passed through a quality control. For the samples processed in Oxford, we PCR amplified five genes from five different chromosomes to give five different sized fragments, and visualized them on an agarose gel (see supplementary Protocol S8 for details). Only samples which successfully displayed at least 4 of the 5 PCR products were used for library preparation and sequencing. Essentially all of the single cell lysates were successfully amplified for DNA and more than 90% of the Oxford samples passed the subsequent quality filter (i.e. quantification by a pico-green assay (Qubit)) before being passed on for DNA sequencing. For samples processed in Eindhoven, a quality check consisting in a PCR of RNase P was performed on some samples. Next, some of the samples were then checked by 1) quantification by Qubit and 2) a test run of sequencing performed at a low number of reads in order to asses the quality of the library before the actual sequencing presented in this paper. Sequencing libraries were successfully prepared from 97% of the samples that passed the initial quality control.

### Contamination

Figure 2A shows the percentage of reads that mapped to the reference genome for 52 cells that were whole genome sequenced. Apart from 7 clear outlier cells (<99% mapped reads), ninety nine percent of the DNA sequences obtained mapped to a reference human genome, indicating that there was effectively no contamination of these samples from non-human sources. This is a significant achievement as there are several published reports of reagent-induced contamination (31–33). In addition, most of the reads are correctly paired, with a low percentage of singletons (0.17 ± 0.06%).

Libraries prepared from single cell DNA could be separated into three main groups according to the level of representation of the human genome: good (9-15 % of non-covered bases), moderate (15–60% of non-covered bases), and bad (90-100 % of non-covered bases). Bad representation of the human genome may be caused by the loss of DNA in the device as supported by the fact that increasing the depth of sequencing for a representative subset of samples did not result in significantly improved coverage. The distribution of the number of reads per cell is given in Figure 2B.

### Coverage

We generated Lorenz plots displaying the fraction of the genome covered versus the fraction of the reads for 39 cells processed in Oxford (Figure 3A-C) and 13 of the single cell sequencing sets obtained from the Eindhoven laboratory (Figure 3D). There is good agreement between these plots and the coverage data shown in Figure 2A. Thus, those plots farthest from the diagonal are for the cells with the poorest whole genome coverage.

We follow Szulwach et al. (34) and calculate *G*, the Gini coefficient for the Lorenz plots, to quantify the uniformity of the genome coverage. The Gini coefficient *G* is calculated as:

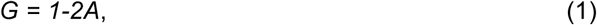

In which *A* is the area under the Lorenz curve. For an ideally uniform coverage of the genome, the Lorenz plot displays a diagonal and the area under the curve is 0.5. *G* = 0 indicates an ideally uniform coverage of the genome. In our study, *G*=0.3 for the sequencing of the bulk of LS174T and many cells have a *G*= 0.5 (see Figure 3D and supplementary Figure S4). In the top tercile of the cells processed in Oxford, corresponding to the highest coverage, *G*= 0.6 ± 0.1 (n=13 cells). For comparison, using commercial instrumentation, Szulwach et al. report *G* = 0.36 ± 0.04 (n=5) for GM12752 cells where the bulk sequencing gives a *G* just below 0.2, but also *G* = 0.6 for another cell type (34).

Thus far most of the single cell sequencing studies only report the coverage results using the Lorenz graph (4,5,14,34). Although the Lorenz graph is effective in reporting which fraction on the genome is not covered, for reporting the distribution of the coverage the so-called coverage graph is more suited and used in (bulk) sequencing experiments. Previously we have reported a coverage graph of single cell sequencing experiments (6). Here, we report a more complete overview of the coverage of our results in Figure 4. The evenness score *E*:

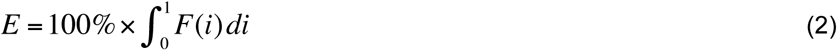

in which *F(i)* is the fraction of the positions with normalized coverage of at least *C(i)/C_ave_* and *C_ave_* is the average coverage provides a metric to quantify the evenness of a read distribution (35). This metric has been has been proposed as one of the 7 metrics to form a description metric for targeted enrichment experiments (36). Here, we use the *E*-score to quantify the evenness of the read distribution in our single cell sequencing experiments. Note that the *E*-score provides a quantitative metric to the coverage graph in the same way as the Gini-coefficient does this for the Lorenz graph. The *E*-score has been put with the coverage and rose from a value 29.3 ± 5.9 % for the poorest results in Figure 4A to around 55-56 ± 10 % for our best results (Fig. 4C.-D). Comparing this result to *E*-scores found in targeted sequencing experiments of 70 ± 5 % (35) and the *E*-score of 87.5% in bulk sequencing (Figure 4D) this suggests that further optimization of the evenness in the read distribution and in the first step of the gDNA amplification process is still needed. Note that the *E*-score is calculated from a normalized coverage distribution. In the SI these normalized coverage graphs are shown in supplementary Figure S6. We show the *E*-score per cell in supplementary figure S7 as well as the coverage graphs in Figure 4, without normalization as this gives a more direct view of the read distribution.

### Allelic dropout

Next, single nucleotide polymorphism (SNP) data were used to obtain estimates of the total recovery of genomic DNA taking into account the near diploid karyotype of LS174T, given that it is mismatch repair defective (Figure 5). From bulk DNA sequencing of both LS174T and LS180 we can easily identify SNPs that are unequivocally heterozygous in both cell lines and which must represent germ line heterozygosity in the patient from whose cancer the two cell lines were derived. Full coverage of the near diploid genome present in LS174T in a single cell sequence would mean, that for such SNPs, both alleles must always be present and observed. If, in the presence of incomplete coverage, we assume that *p* is the probability that one allele of the SNP pair is observed, and that the probability of observing either allele is the same, then *p^2^* is the probability of finding both SNP alleles, *2p(1-p)* is the probability of finding only one of the alleles, and *(1-p)^2^* is the probability that neither allele is found, namely a drop out from both genomes. This is the same binomial result as represented by the frequencies of homozygotes and heterozygotes in a random mating population according to the Hardy-Weinberg law. If *a* is the number of times both alleles are found, *b* the number of times one allele is found and *c* the number of times neither allele is found then the maximum likelihood estimate of *p* is:

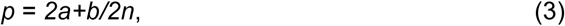

where *n = a + b + c*. This estimate uses the information from all three types of situation rather than just the frequency of heterozygosity, which is a direct estimate of *p^2^* and ignores the information contained in the number of times just one allele is observed. If this calculation is done for a number of SNPs known to be heterozygous in each single cell, then *a*, *b* and *c* can be estimated from the aggregated data on the number of times: *a*, both alleles for the various SNPs are found in the single cell’s DNA sequence, *b*, when only one of the alleles is found and *c*, when neither allele is found. The estimate, *p*, from this aggregate will then be an estimate of the probability of finding any position of the genome in that cell’s DNA sequence once, and *p^2^* will be the probability that the DNA from both the genomes at that site should be present. Thus, assuming the SNPs are a random sample of points on the genome both with respect to position and differential amplification, *p^2^* is an estimate of the true probability of coverage of the total genomic content of the DNA in that singe cell. This is different from the proportion of sequences that map to a reference genome, which does not take into account the presence of two genomes in each cell and so is more or less an estimate of *p* rather than *p^2^*.

Figure 5 shows the results of such an analysis for DNA prepared from the single cells in Oxford using a panel of 12 SNPs known to be heterozygous in the LS174T and LS180 cell lines. The different colours of the vertical bars for each single cell show the proportions of times 2, 1 or no alleles are found, and the cells are ordered from highest to lowest estimate of *p*. The corresponding Lorenz plots for these DNA sequences (Figure 3A-C) show that there is a reasonable relationship between the coverage estimates from the Lorenz plots and the p value estimates. About 44% (17/39) of these single cell DNA sequences give p value estimates of around 0.7 or more, indicating total genomic coverage per cell of around 50%, while about 25% give total coverage of greater than 70%. The overall average p value using the data on all 39 single cells is 0.60 ± 0.25 corresponding to complete coverage of just under 40% (the average p-value for the Eindhoven data in Figure 6 is 0.63 ± 0.22). Out of more than 10,000 reads covering the 13 pairs of alleles for the SNPs, only 63 were ‘incorrect’ in the sense that they were not expected for either allele pair of a given SNP. This indicates a sequencing error rate of less than 1% and also the absence of any contaminating human DNA from external sources, namely other than the cells being analysed.

Additional evidence for the absence of contamination with exogenous DNA was obtained from the density of reads that mapped to male-specific genes on the Y-chromosome, see Supplementary Table S10. Since both the LS174T and the RKO cell lines are derived from female patients and the operators in the Eindhoven laboratory were male, lack of Y chromosome reads provides evidence that there was at least no contamination of Y chromosome reads from them. The male-specific genes used in this analysis are those for which there are no homologies on the X-chromosome as taken from the work of Page and co-workers (37). For almost all male-specific genes we found zero reads mapping to them whereas the mean number of reads per gene on the X-chromosome (taken over all genes listed in the Ensemble human genome annotation GTF file) is over 900 reads/gene on average for these all samples (Supplementary Table S10). This value is to be compared to average number of reads found for the male-specific genes which is 0.4 read/gene (Supplementary Table S10). This effectively rules out exogenous DNA contamination from the male operators in the Eindhoven laboratory to occur and suggests that these reads mapping to male specific genes found corresponds to amplification, sequencing and mapping errors. Since the error rate for sequencing on an Illumina HiSeq system is in the order of 1%. One usually refers to a minimum number of Q30 base (number of base where the error rate is below 1/1000). For 2x125 bp reads of HiSeq, we should have error rate below 1.5% (estimation using an indexed PhiX). The exact value depends on each run. Our data suggest a mapping error of 0.4/900 = 0.04 %.

The heterogeneity of the frequencies of reads (data not shown) between SNPs within single cells suggests dropping out, namely absence of DNA in the initial single cell preparation, as the main reason for lack of complete coverage.

Similarly, Figure 6 gives an estimate of the allelic drop-out for the single cells processed in Eindhoven for which the Lorenz graph is shown in Figure 3D. Note that for the RKO cells, 13 heterozygous SNPs across the genome where used. Again, the results are concordant with the measures of coverage and the Lorenz curves. The poorest cells, with the highest allelic drop-out are right in the far lower corner of the Lorenz plot in Figure 3D, while the best cells, with 60%-70% sharing, correspond to the curves nearest to the diagonal.

In addition to the analysis of single cells from the colorectal cancer-derived cell lines, some single cell whole genome sequences were obtained directly from two fresh colorectal cancers. These were analysed following the same procedure described above using different appropriately chosen sets of SNP markers for each cancer. The results shown in supplementary Figure S5 for a further total of 15 single cells demonstrate that at least comparable quality single cell whole genome DNA sequences can be obtained from fresh tumours as were obtained from the cell line cultures.

Our overall results indicate that the independent analyses of singe cell DNA sequences using two different protocols in different laboratories, but using the same device and instrument, gave comparable results, with perhaps somewhat better coverage using the protocol with alkaline lysis compared to the protocol using proteolytic lysis. Moreover, the results obtained using our valve free devices which are simpler in design and manufacture are comparable with the best published results. For details of the experimental protocols and the use of the instrument see the Methods section and the supplementary protocol S8.

## Discussion

Experimental omics approaches (38, 39) to separately process multiple single cells are increasingly important. Four types of approaches are available for partitioning the molecular contents of one cell from another: 1) Dilution and separately processing; 2) Statistical dilution, tagging and pooling (40), 2) Droplet (41, 42), 3) micro/nanowell (20, 43), 4) Microfluidic trapping (6,7, 44-47).

We found that sequencing genomic DNA extracted from single cells inside our low-cost microfluidic device, gave single cell DNA sequencing results of comparable quality to those reported using more complex and expensive instruments. Our device is valve free and can thus be fabricated by injection moulding a polymer. It is also straightforward and can be operated on a commercial optical microscope or using a custom-built instrument, Cell-O-Matic.

Previously described microfluidic devices isolate single cells using either a physical valve (7) or an oil phase (20) at the time of the lysis and subsequently for the amplification. The use of valves to trap the cells necessitates two-layer devices which are complex and hard to manufacture. By contrast, our devices have no valves and are thus easier to design, manufacture and use. We are able to operate without valves because the liquid flow from different inlets is strictly controlled by air pressure with high accuracy. This, in particular, allows us to exchange reagents in the feeding channel while maintaining the cells trapped until they need to be lysed. The flow rate is minimal as too high flows would dislodge the cells from the traps. The laminar flow conditions in the feeding channel and through the traps ensure that the lysate is pushed through the trap. At a later stage, during the amplification, the solution is confined to the outlet since we can assume there is no significant transport between outlets separated by a channel several centimetres in length. The design of the traps and the mode of lysis and collection of the resulting DNA makes it unlikely that there is significant contamination between the cells trapped on the same chip. Preliminary data obtained by analysis of the LS174T cell line, which is a known mixture of two cell populations (unpublished observations), suggests that there is no major contamination between neighbouring traps on the same chip.

A further proof of the absence of contamination between neighbouring traps can be derived from a subset of data where mRNA was extracted from the captured cells (6). There, PCR of the AXIN2 and Beta-actin genes was used to assess the presence of mRNA in the outlet wells. In those experiments, no mRNA was detected from empty traps adjacent to those where cells were successfully captured and lysed.

Finally, we also consider contamination by exogenous human DNA. The allele analysis of heterozygous SNPs from throughout the entire genome shown in Figure 5 and 6 gives us a good indication that there is no contamination from extraneous human DNA. In addition, we also look at the presence of reads mapped to Y-chromosome genes knowing that the cell lines used in this study are female cell lines. Here, we find generally no reads mapping to those genes (see supplementary Table S10). Some reads do map to a few Y-chromosome genes such as PCDH11Y for all samples but we show that this is due to homologies with genes on the X-chromosome. The density of reads that map to chromosome Y is below 3% and typically 0.05% of the input of a single cell, so should be attributed to amplification and sequencing errors.

When comparing our data to previously published DNA sequencing from single cells we see that the Gini coefficients (see Figure 3) are comparable to results obtained on a commercial system (34). Previously (6) and here we have shown coverage graphs of our single-cell sequencing data. To our knowledge, this has not been done before for single cell sequencing data and we suggest that this should be incorporated in future single sequencing experiments as this gives a better insight in the read distribution in these experiments and to what extent reliable SNP calling can be performed. Finally, we have presented our results in terms of maximum likelihood estimate of allelic dropout p and find this value to be 0.6 ± 0.2.

The first commercial microfluidic device method for processing single cells for sequencing (17) was known to suffer significantly from the capture of doublets rather than single cells. Our approach has the advantage that we can take an image of trapped cells to confirm the single cell occupancy of each trap before proceeding to sequencing.

The more recently emerging droplet-based single cell fluidics and dilution tagging and pooling approaches offer the highest throughput (1-10,000s of cells) compared to 10-100s of the microfluidic trapping approaches. However, an advantage of our approach, is that it can be used to extract and process RNA from the same cell as the DNA (6); such multi-omic characterization will be important for making the connection between genotype and molecular phenotype to gain a better understanding of cellular mechanisms and to better select the mutations that may be driving a cancer phenotype and which might be candidates for targeted therapy.

For such integrative omics applications it is important to know that the comparative performance metrics of our single cell processing devices are equivalent to other types of devices and approaches. We can conclude that our DNA sequencing results show that the output of our device is at least comparable to, if not better than the valve-based commercial devices and offers advantages over non-microfluidic approaches such as a very low contamination level.

## Availability

### Accession Numbers

Our single cell sequencing data is available as.bam files at the SRA: (https://www.ebi.ac.uk/ena/browse) under accession numbers ERS2168085 - ERS2168123.

### Supplementary Data

Supplementary Data are available at NAR online.

## Acknowledgements

The authors gratefully acknowledge funding from the European Commission under the Seventh Framework Programme (FP7/2007–2013) under grant agreements number 278204 (Cell-O-Matic).

